# The reward positivity is insensitive to reinforcer devaluation

**DOI:** 10.1101/2025.03.27.645774

**Authors:** Lindsay S. Shaffer, Holly D. Crowder, Peter A. Kakalec, Lam T. Duong, Craig G. McDonald, James C. Thompson

## Abstract

Successful behavioral adaptation requires an ongoing assessment of rewarding outcomes based on one’s current state. A frontocentral ERP associated with reward feedback, the reward positivity (RewP), has been linked to reflect information about reward value and motivational states. It is, however, unclear if changes in the RewP are influenced by changes in reward value as a function of motivational state. To examine this, hungry participants (n=31) completed two rounds of a modified Doors Task incorporating Pavlovian conditioning during EEG recordings and obtained feedback associated with sweet and savory food reinforcers equally matched in pleasantness and desirability. Participants underwent reinforcer devaluation, a paradigm designed to isolate inference-based behavior based on decreasing reward value, in between rounds by eating one of the foods to satiety. Prior to devaluation, participants were hungry and rated both food reinforcers equally pleasant. After devaluation, participants were sated and rated the devalued food, but not the non-devalued food, significantly less pleasant, suggesting a sensory-specific change in reward value. Logistic regression of win-stay/lose-switch behavior during the Doors Task shows participants made sensory-specific adjustments in food preferences during post-devaluation. Non-parametric permutation tests based on the tmax statistic performed revealed no significant differences in RewP amplitudes, suggesting the RewP is insensitive to reinforcer devaluation. This could not be explained by differences in perceived pleasantness or desirability. These findings suggest that affective and motivational factors such as tracking inferences based on decreases in reward value did not modulate the RewP.

## Introduction

A lot of decisions we make are based on what we likely expect in the future. Being able to flexibly change your behavior due to adjustments in those expectations, however, is essential to survive in a constantly changing environment. One way adjusting expectations could occur is due to changes in how we perceive expected rewarding outcomes. For example, one could consume their favorite meal for lunch but no longer desire their favorite meal for dinner because what was once rewarding is no longer such due to consuming your favorite meal to satiety. One outstanding question in the field of cognitive neuroscience is whether changes in expected rewarding outcomes reflect direct experience (e.g., going to the same restaurant for dinner) or inference (e.g., going to a different restaurant for dinner but same cuisine). While a lot of decisions we make represent the combination of both kinds of expectations (Feher da Silva & Hare, 2020; Kahnt & Schoenbaum, 2021), disassociating behavior exclusive to inferred or direct experiences is needed to understand how different kinds of expectations contribute to behavior and how these processes are reflected in the mammalian brain.

One approach to examine whether behavioral adjustments are guided by inference is the reinforcer devaluation paradigm. Used in both Pavlovian (Holland & Rescorla, 1975; Holland & Straub, 1979) and instrumental (Adams & Dickinson, 1981; Colwill & Rescorla, 1985b; Colwill & Rescorla, 1990) conditioning, participants receive training with more than one reinforcer to learn the association between a stimulus or action leads to a specific outcome. Using multiple reinforcers allows the opportunity to utilize within-participants design and show behavior is influenced by the sensory properties of one reinforcer as opposed to its affective or general properties (O’Doherty, 2014). After learning the associations between cues or actions with outcomes, the value of one of those reinforcers is decreased in the absence of cues or responses associated with the reinforcer outside of the experiment context. Decreasing the value of the reinforcer can be accomplished by having the participant consume the reinforcer until they reach satiation. Following devaluation, participants complete the same task as they did prior to devaluation except the stimuli or actions are presented in extinction, or the omission of the reinforcer. Since post-devaluation testing is conducted during extinction, reduced responding toward stimuli associated with devalued outcomes is not easily explained by direct experience, suggesting that the current value of the reinforcer was inferred in the absence of that reinforcer’s presence in order to adjust behavior.

Aside from its utility as an assay of goal-directed behavior, reinforcer devaluation effects have broader implications. Specifically, devaluation effects reflect what kind of information is task-relevant. According to Pavlovian learning theories, two forms of Pavlovian predictions can co-occur: predictions based on associations between the stimulus and sensory-specific properties—the identity of the reinforcer independent of its metabolic properties (e.g., pretzels, milk chocolate)—and predictions based on associations between the stimulus and its motivational or affective properties (e.g., tasty snacks; Konorski, 1967). It’s thought that the associations between stimuli and the sensory-specific properties of reinforcers are what underlie the construction and updating of a cognitive map of the current task space (Costa et al., 2023; Wilson et al., 2014). This is because in order for agents to successfully learn the associations between stimuli and their outcomes, agents need to be able to represent task-relevant information correctly. Thus, in order for specific adjustments in behavior to occur, predictions about expected outcomes must be specific and beyond their value. While specific brain regions such as the orbitofrontal cortex and basolateral amygdala have been implicated in sensory-specific representations (Howard et al., 2020; Gottfried et al., 2003), utilizing electrophysiological methods such as scalp electroencephalography (EEG) could provide insight into the dynamics of information processing associated with state representations.

Although few EEG studies have utilized reinforcer devaluation, those that have nonetheless provided valuable insight into the temporal resolution underlying perceived reward value. In one study using verbal instructions as a devaluation technique and instrumental conditioning, Luque and colleagues (2017) demonstrated greater P1 amplitudes toward devalued reinforcers following devaluation compared to non-devalued reinforcers in occipital electrodes, whereas greater P3b amplitudes were observed for non-devalued reinforcers compared to devalued reinforcers following devaluation in occipital electrodes. These results would suggest the co-occurrence of goal-directed and habitual systems in visual cortices relevant for memory recall and subsequent context-updating. In another study, Huvermann and colleagues (2021) had participants complete two separate sessions of a Doors Task in which participants could win a highly preferred snack, a moderately preferred snack, or a low preferred snack. Crucially, participants completed the Doors Task either without eating any food or eating their highly preferred snack. When viewing pictures of devalued food, participants exhibited modulated P3 amplitudes, suggesting that the P3 component is sensitive to changes in subjective preferences. Together, both studies highlight how different ERP components encode rewarding experiences relevant for adaptive behavior.

One event-related potential (ERP) implicated in reward processing is the reward positivity (RewP). The RewP observed at frontocentral regions of the scalp approximately 200-350 ms following feedback associated with reward receipt (Proudfit, 2015). Initially under the RL framework, the RewP was proposed as a signal reflecting prediction errors facilitated by midbrain dopaminergic projections to the prefrontal cortex as unexpected rewards elicit greater RewP amplitudes than expected rewards (Holyrod & Coles, 2002; Holroyd & Yeung, 2012). Recently, it’s been shown that the RewP is sensitive to hedonic liking even when controlling for the unexpectedness of the feedback delivery (Brown et al, 2022; Huvermann et al., 2021; Peterburs et al., 2019). This would suggest that computations underlying the RewP might not participate in a unitary process of reward processing, but that reflect multiple dimensions of liking, wanting, and learning of rewarding experiences.

Interestingly, recent work on examining the RewP’s sensitivity to subjective preferences and motivational state implies a potential role in encoding state representations. For example, RewP amplitudes were greater for highly preferred rewards versus less preferred rewards (Peterburs et al., 2019) and self-reported measures of pleasantness were correlated with RewP amplitudes in response to preferred reinforcers (Brown et al., 2022). Both of these reports support the RewP sensitive to hedonic liking through reward preferences. In addition to support for the role of the RewP in encoding reward value that’s contextual to the participant’s preferences, recent work has demonstrated the RewP showing contextual sensitivity to reward value as it relates to factors associated with affective or motivational state. For example, deprivation of rewards relevant to nicotine withdrawal lead to increasing the value of those rewards as indexed by the RewP (Baker et al., 2016). Additionally, hunger levels were associated with larger RewP amplitudes when viewing food rewards but not money rewards but RewP amplitudes for both food and money rewards were similar when participants were sated upon viewing feedback (Banica et al., 2023). These findings suggest that motivational state modulates reward value at the time of feedback receipt and this information is contributing to the processes underlying the RewP. Although these reports suggest a contextual role of motivational state contributing to the RewP, it’s unclear how sensory-specific representations attribute to participants’ motivational states by consuming several different foods at a standardized time point in the experiment. Therefore, an outstanding question is whether the RewP reflects reward value as a function of current motivational state because this form of sensitivity would suggest the encoding of task-relevant information conveyed by state representations.

We sought to examine the RewP’s sensitivity to reward signaling as a function of motivational state by implementing a modified version of the Doors Task requiring participants to learn relationships between feedback and outcomes (Gottfried et al., 2003; Proudfit, 2015; Pool et al., 2019). Participants completed two rounds of our modified Doors Task to win food rewards and eating one of those food rewards between rounds in order to devalue one of the foods. Although most RewP studies use secondary reinforcers such as points or money, we opted to incorporate primary reinforcers such as food as hunger is a naturally occurring state. It’s also been shown that how food rewards are presented does not constrain how its hedonic impact is perceived. For example, hungry humans and animals find food rewards pleasant regardless if the food was presented visually (O’Doherty, 2002), as odors (Cabanac, 1971; Cabanac & Duclaux, 1970; Howard & Kahnt, 2017; Small et al., 2001; Gottfried et al., 2003), and through taste (Rolls et al., 1981; Rolls et al., 1983). This could be because we use different kinds of sensory information across modalities to create representations relevant for food-seeking behavior. Moreover, updating information in one modality might transfer to how we perceive the same food reward in a different modality. Thus, featuring food rewards affords a unique opportunity to generalize our RewP results because food reward involve multi-modal flexibility in how representations are created, maintained, and updated.

Since we incorporated two different yet equally pleasant and desirable food rewards, we should expect to evoke a RewP during the pre-satiety session. Moreover, reward feedback waveforms comprising the RewP should show similar amplitudes during the pre-satiety session. If the RewP is sensitive to changes in motivational states and reward value, then we predict that the RewP amplitude would significantly decrease during post-satiety. For the P1, we predict greater amplitudes in response to devalued feedback compared to non-devalued feedback during the post-satiety session, whereas amplitudes for both raw reward feedback would evoke similar P1 amplitudes during the pre-satiety session.

## Materials and Methods

### Participants and apparatus

Due to the novelty of this study, we did not perform an *a priori* power analysis. Instead, we determined our sample size based off of previous studies that have reliably evoked a reward positivity. Most of the sample sizes from these studies ranged between 20-40 participants. Thus, our goal was to have a total sample size of at least 30 participants. Forty-one healthy volunteers consented to enroll in this experiment and received compensation of either course credit or $30.00 upon completion of all study procedures. Participants were recruited from the George Mason University Department of Psychology participant recruitment pool. All participants reported normal or corrected-to-normal vision. Seven participants were excluded due to equipment issues during the recording sessions, but behavioral data for two participants from this subgroup were acquired. Three participants did not have EEG recordings completed as the equipment was unable to accommodate specific hairstyles (e.g., box braids, dreadlocks) but still completed the behavioral components of this study. Although behavioral data from thirty-six participants were recorded successfully, four participants were missing pleasantness rating scores and two participants were missing hunger rating scores. The final data set composed of thirty-one participants for the EEG data analysis, thirty-two participants for the pleasantness and desirability scores analyses, and thirty-four participants for the hunger score analysis (see Table 1 for demographics). The study was approved by the George Mason University International Review Board and in accordance with guidelines from the Declaration of Helsinki.

**Table 1.**
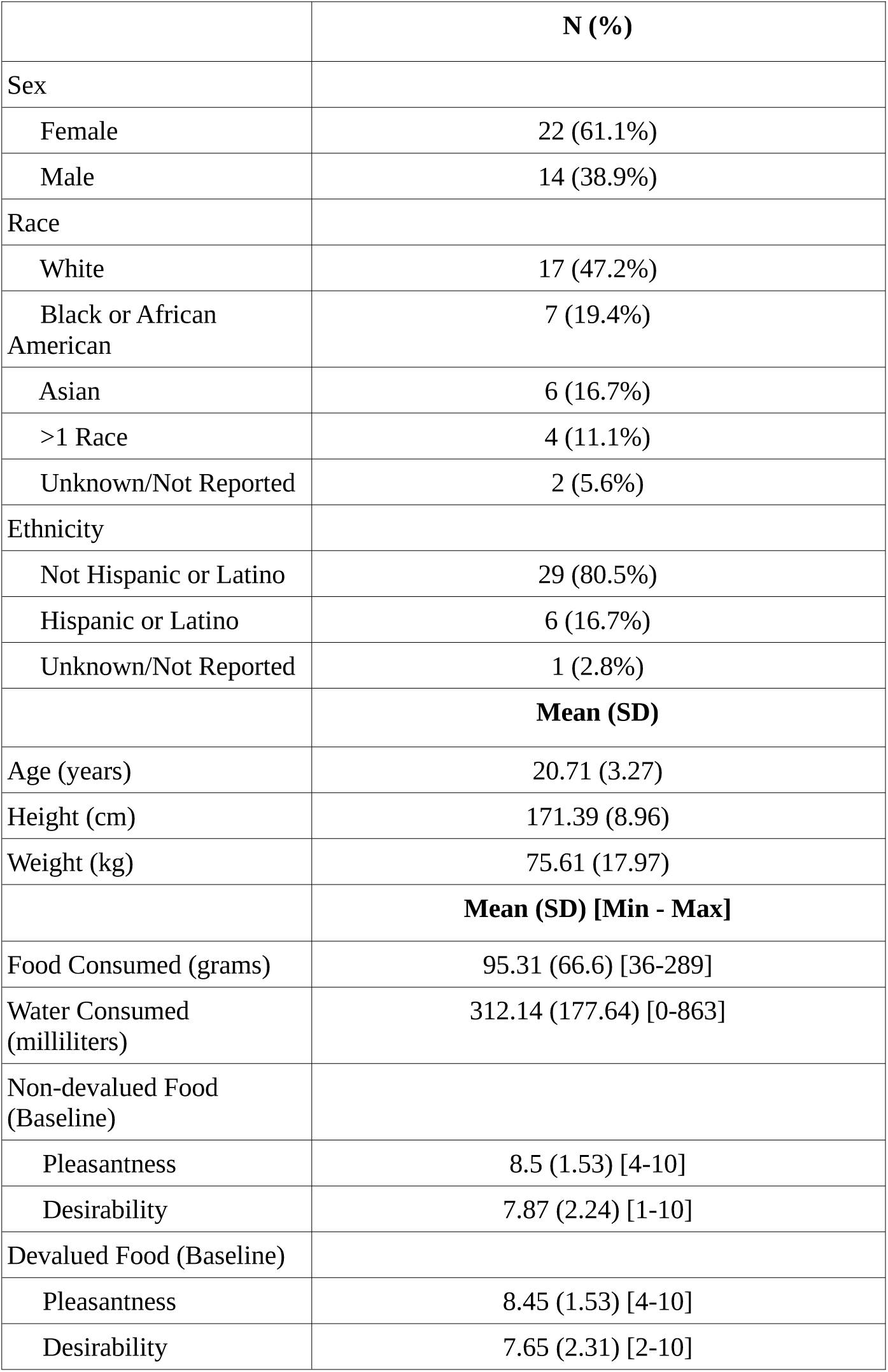
Participant demographics (n=36).

|  | N (%) |
| --- | --- |
| Sex |  |
| Female | 22 (61.1%) |
| Male | 14 (38.9%) |
| Race |  |
| White | 17 (47.2%) |
| Black or African American | 7 (19.4%) |
| Asian | 6 (16.7%) |
| >1 Race | 4 (11.1%) |
| Unknown/Not Reported | 2 (5.6%) |
| Ethnicity |  |
| Not Hispanic or Latino | 29 (80.5%) |
| Hispanic or Latino | 6 (16.7%) |
| Unknown/Not Reported | 1 (2.8%) |
|  | **Mean (SD)** |
| Age (years) | 20.71 (3.27) |
| Height (cm) | 171.39 (8.96) |
| Weight (kg) | 75.61 (17.97) |
|  | **Mean (SD) [Min - Max]** |
| Food Consumed (grams) | 95.31 (66.6) [36-289] |
| Water Consumed (milliliters) | 312.14 (177.64) [0-863] |
| Non-devalued Food (Baseline) |  |
| Pleasantness | 8.5 (1.53) [4-10] |
| Desirability | 7.87 (2.24) [1-10] |
| Devalued Food (Baseline) |  |
| Pleasantness | 8.45 (1.53) [4-10] |
| Desirability | 7.65 (2.31) [2-10] |

**Table 2.** Self-report measures before and after reinforcer devaluation.

|  | **Pre-Satiety**<br>**Mean (SD) [Min - Max]** | **Post-Satiety**<br>**Mean (SD) [Min - Max]** |
| --- | --- | --- |
| Hunger<sup>1</sup> | 8.20 (1.12) [4 - 10] | 3.26 (1.81) [1 - 6] |
| Pleasantness<sup>2</sup> |  |  |
| Non-devalued food | 8.31 (1.51) [3 - 10] | 7.43 (1.66) [3 - 10] |
| Devalued food | 8.46 (1.50) [5 - 10] | 5.65 (2.54) [1 - 10] |
| Desirability |  |  |
| Non-devalued food | 8.00 (1.86) [2 - 10] | 5.75 (2.72) [1 - 10] |
| Devalued food | 8.12 (1.87) [4 - 10] | 2.81 (1.76) [1 - 7] |

<sup>1</sup> Hunger scores were collected from 34 participants.
<sup>2</sup> Pleasantness and desirability scores were collected from 32 participants.

Participants were test individually within a cubicle using a standard Windows PC. Stimulus presentation was programmed in Python 3.6 and incorporated the PsychoPy library (Pierce et al., 2014). Responses were made on a standard PC keyboard. Since data collection occurred during the COVID-19 pandemic, experimenters implemented guidelines for practicing EEG in compliance with the university’s practices along with practices as described in the literature for EEG research (Simmons & Luck, 2020). Participants were required to abstain from eating any food or drink any liquids other than water for at least three hours prior to their appointment time. As majority of participant appointments were completed in the morning, participants typically reported not eating or drinking anything since the night prior to their appointment.

#### Experimental procedure

Following informed consent, participants completed a demographic survey (see Table 1). After completing the demographics survey, participants were presented another survey which listed 14 snack foods that were evenly divided into savory and sweet categories which they subsequently rated each food item’s pleasantness and desirability on a 10-point Likert type scale. The foods used were described in a previous study (Reber et al., 2017). The savory foods consisted of Cheetos^®^, Fritos corn chips, Goldfish^®^ cheddar snacks, low-sodium Plantar’s whole cashews, Skinny Pop non-cheddar white popcorn, pretzels, and Ritz Bits cheese crackers. The sweet foods consisted of Haribo gummy bears, M&M’s^®^, mini marshmallows, Oreos, Reese’s peanut butter cups, Skittles, and sour gummy worms. After the participant completed the food ratings, the experimenters calculated a preference score as the sum of the pleasantness and desirability ratings for each food item and followed a forced choice determination as described in Reber et al. (2017). This resulted in the participant choosing a food in one category (e.g., sweet) that was comparable in pleasantness and desirability ratings to the other chosen food in the other category (e.g., savory). After the participant completed the food preference survey and choice determination, the experimenters prepared the participant for EEG data collection (for details, see EEG acquisition).

Once participants completed the food preference survey and preparing for EEG data acquisition, they began a modified version of the Doors Task (Figures 1A, 1B). Participants were presented with instructions on completing the task, such that they should try to pick doors to win as many food prizes as possible. Participants were informed that sometimes they might win a food prize or they might not win anything at all and to use any type of strategy they want to win. After verifying participants understand the instructions as assessed by the experimenter, participants were presented with three identical doors on a black screen. Participants could respond to each stimulus at their own pace by pressing either the “A” key for the left door, the “S” key for the middle door, and the “D” key for the right door. All participants used their left hand on the keyboard for responses. After a response was made, a jittering inter-stimulus interval (ISI) was presented in the form of a fixation cross between 700-1000 ms. After the ISI, the participant were presented with feedback positioned at the center of the screen for 1000 ms. The stimuli used for feedback were adjusted for luminosity. Participants either saw a magenta or blue arrow if they won either type of food and saw a zero if they did not win any food for that round. The color of the arrow for each type of food was counterbalanced across participants. Following feedback, a picture of the outcome associated with the feedback would be presented directly above the feedback itself for 1500 ms. In this context, the specific savory or sweet food item chosen by the participant would be presented above feedback associated with food prizes, whereas a picture of an empty plate was associated with not winning any food. Despite feedback presentation occurring independent of door selection, concurrently presenting the feedback and outcome irrespective of the actions taken facilitated the Pavlovian association between feedback (e.g., arrows, zero) and outcomes (e.g., food, empty plate pictures). Outcomes were also presented directly above the feedback in order to minimize ocular movements from the participant. Following outcome presentation, a jittering inter-trial interval (ITI) in the form of a fixation cross was presented for 700-1500 ms. Following the ITI, participants viewed instructions that prompted “Press the space bar to begin the next round.” This allowed participants to combat any potential fatigue and complete each trial at their own pace. To further combat participant fatigue, participants were required to rest for a minimum of 30 seconds prior to the beginning of the next block. The entire modified Doors Task consisted of 240 trials (10 blocks of 24 trials) in which participants received no rewards 50% of the time, the devalued feedback 25% of the time, and the non-devalued feedback 25% of the time. The outcome schedule of the modified Doors Task maintained an equally probabilistic schedule of devalued and non-devalued feedback types while allowing the learning of specific rewarding outcomes. The outcome schedule of the modified Doors Task also ensured that rewards and losses were presented in an equally probabilistic manner, as components such as the N2 which has spatiotemporal overlaps with the RewP can be influenced by novelty effects (Brown et al., 2022).

**Figure 1.**
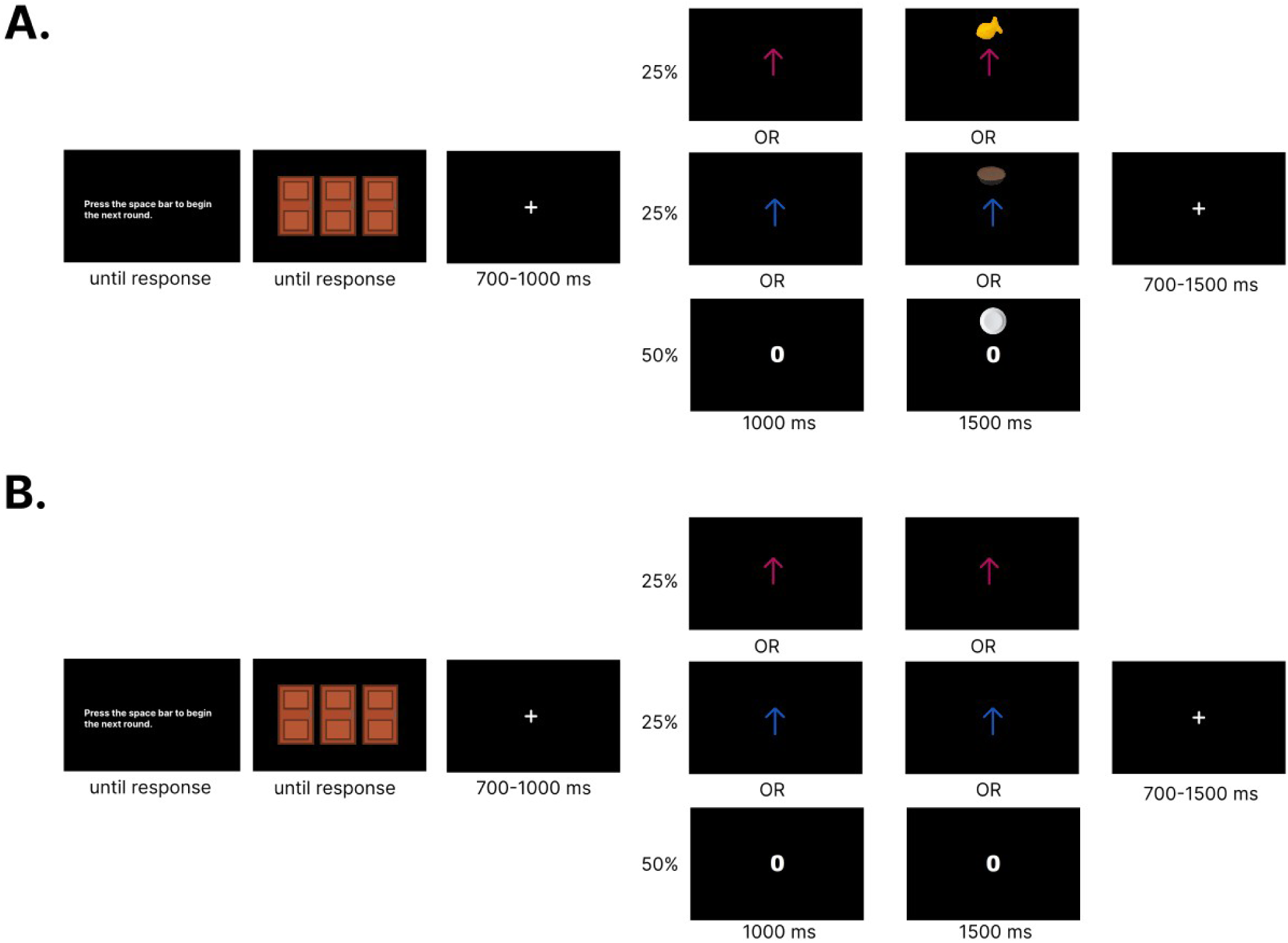
Experimental paradigm. a) Trial schematic during the conditioning (pre-satiety) session, and b) extinction (post-satiety) session. Food rewards were associated with either a blue or magenta up arrow. No reward was associated with a zero with a picture of an empty plate. Designation to which color arrow was assigned to each food reward was counterbalanced.

**Figure 2.**
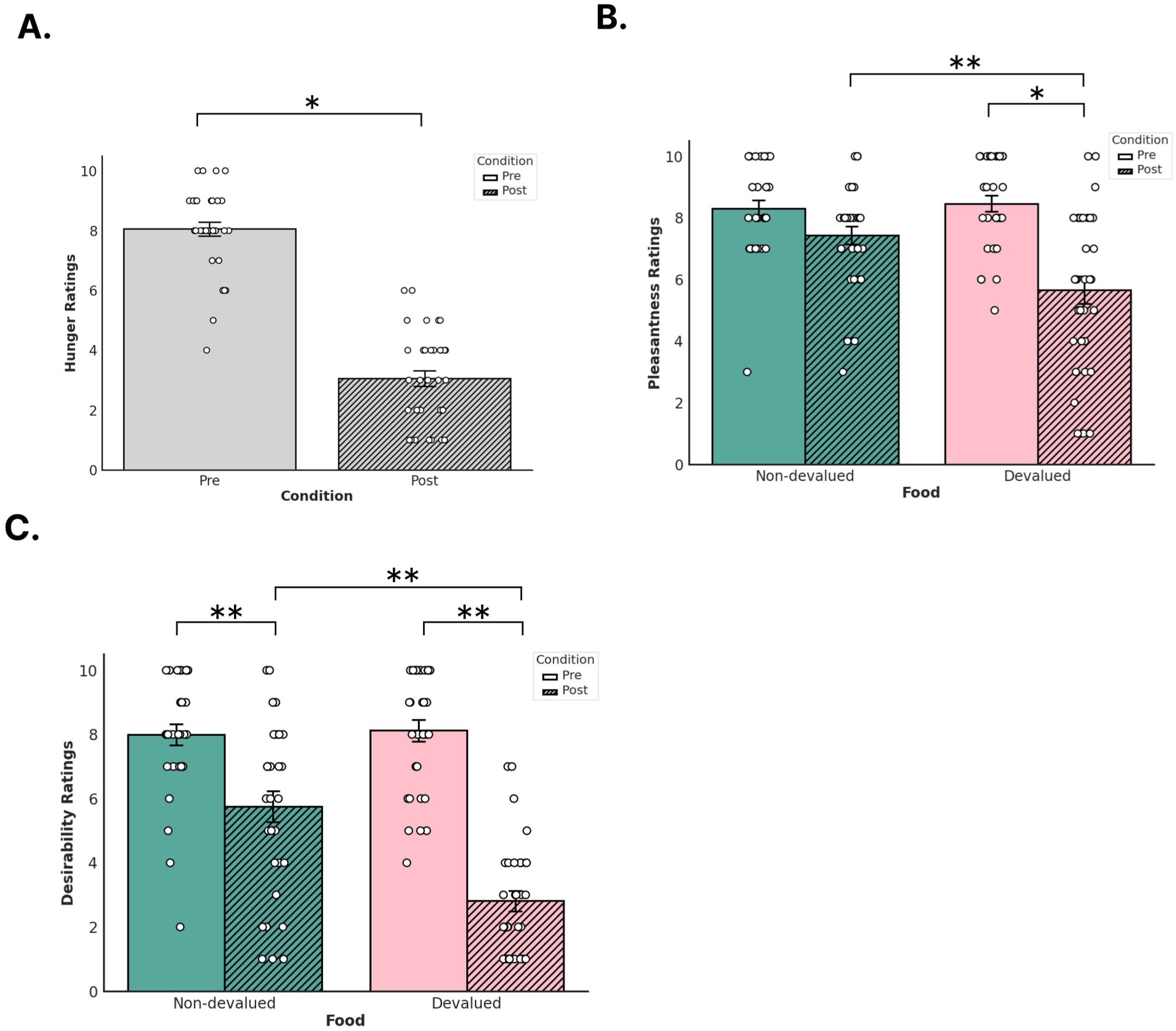
Self-reported measures before and after devaluation. Participants provided ratings for a) hunger, b) pleasantness for non-devalued (orange) and devalued (green) reinforcers, and c) desirability for non-devalued and devalued reinforcers. Solid colored bars denote pre-devaluation and striped bars denote post-devaluation. Error bars denote standard error (SE). White dots denote single data points from participants.

After participants completed the Doors Task during the pre-satiety session, the experimenter asked the participant how hungry they current are, how much they liked each food, and how much they wanted each type of food. Responses for each question were reported on a 10-point Likert-type scale. After responses were recorded, the experimenters left the testing room to measure food out of sight from the participant. The food chosen as the devalued outcome was determined based on a stratified randomization schedule that divided participants into subgroups based on reported sex (e.g., male, female) and which type of food was consumed (e.g., salty, sweet; see Kernan et al., 1999). Food and water was measured using a scale and documented prior to providing the participant with the food. Following measuring the food, the experimenters presented the food and gave instructions to the participant to eat as much food as they wanted until they were physically incapable of eating the food. The experimenters left the room as participants ate the food. After the participants notified the experimenters that they were done consuming food, the experimenters removed the food and water to measure. While one experimenter measured the remaining food and water, the other experimenter asked participants how hungry they were, how much they liked a certain food, and how much they wanted a certain food in that moment using the same 10-point Likert-type scales prior to reinforcer devaluation. After obtaining responses from participants, the experimenters began the post-satiety session in which participants completed the identical Doors Task as before but conducted in extinction (i.e., feedback without outcomes were presented). The experimenters instructed participants that their results would still be recorded despite the food or plate pictures no longer being shown. Experimenters also instructed participants to win as many food prizes as possible as this will determine what participants will eat after the task is completed, based on instructions from Morris et al. (2015). After the completion of the second Doors Task, participants were debriefed by the experimenters and compensated for completing the study.

### Data acquisition

#### EEG acquisition

EEG data were collected using a Brain Vision actiChamp amplifier and Brain Vision Recorder version 2.1 software (Brain Products, Inc.). EEG data were recorded using a 32 actiCAP electrodes placed on the head using a fitted cap calibrated according to the 10-20 system (American Electroencephalographic Society, 1994). An in-cap ground electrode was positioned anterior to channel Fz. Channel Cz was used as the online reference for the other 31 electrodes. Since channel Cz is central to test our hypothesis, data for channel Cz were recovered offline. Impedance from electrodes were kept below 25 kΩ prior to the start of the experiment. The recordings were digitized at a sampling rate of 500 Hz and a low- and high-pass filter were applied to signals during online recording between 0.1 Hz and 249 Hz.

#### Data pre-processing

EEG pre-processing was completed in MATLAB R2021a (The MathWorks; https://www.mathworks.com/). Initial steps were conducted using EEGLAB 2020.0 and ERPLAB 10.1 (Delorme and Makeig, 2004; Lopez-Calderon and Luck, 2014). Data were re-referenced to the averaged left and right mastoids (TP9, TP10). A bandpass filter at 0.1-30 Hz was applied to the data and then subsequently downsampled to 250 Hz. Data were segmented into epochs with a duration of 1000 ms ranging from -200 ms before stimulus onset to 800 ms post-stimulus onset during the following events: presentation of the doors, presentation of the feedback, and presentation of the outcome. A baseline correction was then applied to all epochs to 200 ms prior to feedback onset to 0 ms. Data were then visually inspected and electrodes with a high amount of noise in their signals for at least 20% of the epochs were removed. Following the removal of bad channels, epochs were rejected if they had voltages greater than 100 mV between sample points or a voltage difference of a minimum of 200 mV per segment. The average of number of trials was as follows: pre-satiety non-devalued feedback (M = 57.96; SD = 7.75), pre-satiety devalued feedback (M = 58.16; SD = 6.85), pre-satiety no win feedback (M = 115.74; SD = 15.12), post-satiety non-devalued feedback (M = 59.32; SD = 1.44), post-satiety devalued feedback (M = 59.03; SD = 1.25), and post-satiety no win feedback (M = 118.19; SD = 2.57). The data for each participant’s pre-satiety and post-satiety sessions were concatenated and decomposed using independent component analysis (ICA). Concatenating data for both sessions resulted in identical ICA weights for each participant’s pre- and post-satiety recordings. ICA components corresponding to eye blinks, saccades, or signal drop-out were removed (M = 5.6 components removed; SD = 1.77). After the removal of ICA components, bad channels that were removed prior were interpolated. Following interpolation, an averaged ERP waveform was created using the *pop_averager()* function in ERPLAB (Lopez-Calderon & Luck, 2014).

For our analyses, we quantified the time window based on previous suggestions in the literature on identifying RewP time windows based on differences across experimental paradigms (Sambrook & Goslin, 2015). Although the RewP is reported to be evoked between 240-340 ms post-feedback, variations in time windows used are experiment-dependent and influenced by variations in the N2 and P3 component (Sambrook & Goslin, 2015; Krigolson, 2018). In order to account for this variation, we determined the average most negative trough measured between 240-340 ms post-feedback using a grand average of non-devalued feedback and devalued feedback for all participants collapsed during the pre-satiety session. We reasoned that the most parsimonious explanation of the RewP is that it manifests as a modulated N2 component (Krigolson, 2018). The average peak latency of the most negative trough at 316 ms was identified and a time window of 291-341 ms was centered on this latency. After creating our time window, we extracted the mean amplitude recorded at 291-341 ms post-feedback for Fz, Cz, and Pz. We reasoned that although the RewP has been reported to be maximal evoked in medial frontal regions such as channel FCz, maximal amplitudes have also been reported in Fz, Cz, and posterior regions corresponding to channel Pz (Sambrook & Goslin, 2015; Cavanagh, 2015; Krigolson, 2018). As the latency of the RewP can vary within and across experimental designs, we extracted mean amplitudes in two preceding time windows corresponding to 189-239 ms and 240-290 ms in order to determine whether the RewP occurs outside of our initial time window of 291-341 ms. Since we sought to replicate results from a previous ERP study with reinforcer devaluation by measuring the P1 (Luque et al., 2017), we identified an average peak latency from the grand average of 126 ms by measuring between 60-200 ms at channel Oz post-feedback which resulted in a 50 ms time window of 101-151 ms centered on the peak latency. Once our time window was identified, we extracted the mean amplitude recorded at 101-151 ms post-feedback for channel Oz.

#### Data analysis

For the behavioral data, we calculated the mean scores of perceived pleasantness and perceived desirability for the devalued and non-devalued food at pre-satiety and post-satiety. We also calculated the mean hunger score reported by participants at pre-satiety and post-satiety. Self-report measures of perceived pleasantness, desirability, and hunger have been used in human reinforcer devaluation studies to confirm the presence of manipulation effects (Pool et al., 2019; Reber et al., 2017; Howard & Kahnt, 2017; Gottfried et al., 2003). To account for individual variation with pleasantness and desirability scores, we performed a linear mixed-effects model using the *nlme* library in R (Pinheiro et al., 2021; R Core Team, 2013). By using a linear mixed-effects model, this allowed us to account for the nested structure of our data in which participants varied in self-report ratings across both conditions and food types. Specifically, we analyzed whether perceived pleasantness or desirability scores were influenced by the interaction of our fixed effects predictors of condition (pre vs post) and food type (devalued and non-devalued). Similarly with our models for perceived pleasantness and desirability scores, a linear mixed effects analysis was performed to determine whether hunger scores were influenced by the fixed effect predictor of condition (pre vs post).

We also examined the likelihood of adjusting behavioral strategies as a function of what type of feedback participants received. Previous studies on reinforcer devaluation have examined averaged response rates or autonomic arousal toward non-devalued and devalued reinforcers (Allman et al., 2010; Pool et al., 2019; Thompson et al., pre-print; Howard & Kahnt, 2017; Gottfried et al., 2003), which could not be measured using gambling tasks such as the Doors Task. Previous gambling tasks, however, have quantified the likelihood of switching responses as a function of the previous trial outcome (Hayden et al., 2008; Hayden et al., 2011). For this analysis, we did not have specific *a priori* predictions regarding how switching behavior would be influenced by reinforcer devaluation as no previous reinforcer devaluation study has examined response rates in this manner. We reasoned, however, that if non-devalued feedback was considered more pleasant than devalued cues following devaluation, then it’s likely that participants would indicate their preference for non-devalued feedback based on their behavior in our task. To determine how feedback influenced subsequent choice behavior, we performed a binomial logistic regression using a mixed-effects model with feedback and session as predictors and whether responses were labeled as a switch as our outcome variable using the *stats* package in R and custom scripts in Python (R Core Team, 2013). Responses were labeled as a switch if the current trial’s key press upon selecting a door did not match the key press for the previous trial, whereas responses were labeled as staying if the current trial’s key press matched the previous trial’s key press. Therefore, responses closer to a 0 suggest participants were more likely to repeat the same choice as they did for the previous trial, whereas responses closer to a 1 suggest participants were more likely to make a different choice from the one taken from the previous trial. For our model, we used No Reward during the pre-satiety session as our reference point testing for the main and interaction effects of condition (pre vs post) and feedback type (devalued, non-devalued, and no reward).

Prior to performing statistical analyses, we calculated several difference waveforms as our RewP. We calculated a Non-devalued RewP (“Nondev RewP”) as the difference between non-devalued feedback and no wins and calculated a Devalued RewP (“Deval RewP”) as the difference between devalued feedback and no wins (Figures 5A, 5B). Both the Nondev and Deval RewPs were primarily observed over posterior regions along the scalp and thus were plotted at channel Pz (Figures 5A, 5B). To detect reliable differences in whether the mean amplitudes recorded for our ERPs were influenced by either condition (pre vs post), feedback type (Nondev RewP vs Deval RewP), or the interaction between condition and feedback type, we performed permutations using a two-samples t-test. Since we are interested in observing the effects of brain activity due to the interaction between feedback and condition, we performed the non-parametric permutations tests using a grand average difference wave by taking the difference between the sessions by subtracting the pre-satiety and post-satiety sessions for the Nondev RewP and Deval RewP (i.e., taking the difference of a difference for each RewP). By comparing the grand average Nondev RewP with the grand average Deval RewP, our permutation test assumes no differences between Nondev RewP and Deval RewP regardless of condition. For the P1, we compared the grand average non-devalued feedback and grand average devalued feedback measured at channel Oz. After the initial t-tests were performed, t-tests were performed again 1000 times with data shuffled within each participant. All permutations tests performed for each channel had the same number of epochs. This is because of the possibility that any potential spatiotemporal correlations between adjacent EEG electrodes and the unknown modulation effects of reinforcer devaluation on ERPs might increase the likelihood of type I errors in our results. Since permutation testing could increase the likelihood of type I errors, we corrected p-values from multiple comparisons by calculating the proportion of the maxT statistic that is greater than or equal to the critical t-statistic and divided by the number of permutations (Holmes et al., 1996; Nichols & Holmes, 2001). Since we had multiple time windows, we performed each permutation test for each of our different time windows (P1 and RewP) using the steps above.

## Results

### Sensory-specific reinforcer devaluation alters levels of hunger and self-reported food preferences

Linear mixed-effects models showed that satiety (t(33) = 13.32, beta = 4.91, p = 0.001 [CI 95%: 4.16, 5.66]) significantly decreased after reinforcer devaluation compared to before suggesting that participants reported feeling sated after eating food. Our linear mixed-effects models measuring self-reported ratings of food pleasantness showed a significant interaction effect between food type and condition (t(93) = -3.19, beta = -1.94, p = 0.002 [CI 95%: -3.14, -0.73]) and a main effect for condition (t(93) = -2.04, beta = -0.87, p = 0.044 [CI 95%: -1.73, -0.02]). A main effect for reward type was not significant (t(93) = 0.36, beta = 0.16, p = 0.716 [CI 95%: -0.70, 1.01]). In this case, both devalued and non-devalued reinforcers were similarly rated in their perceived pleasantness prior to devaluation. After devaluation, however, the pleasantness rating of the devalued food reinforcer significantly decreased whereas the pleasantness rating for the non-devalued reinforcer remained similar prior to devaluation. Our linear mixed-effects models measuring self-reported ratings of food desirability showed a significant interaction effect between food type and condition (t(93) = -4.72, beta = -3.06, p < 0.001 [CI 95%: -0.78, 1.03]) and a main effect for condition (t(93) = -4.91, beta = -2.25, p < 0.001 [CI 95%: - 3.15, -1.34]). A main effect for reward type was not significant (t(93) = 0.27, beta = 0.12, p = 0.785 [CI 95%: -4.34, -1.77]).

Since we found interaction effects between food type and condition for our pleasantness and desirability ratings, we calculated the estimated marginal means (EMMs) of our linear mixed-effects models of pleasantness and desirability separately using the *emmeans* package in R (Lenth, 2024). Calculating EMMs can help determine whether any pairwise comparisons between the means of each factor are statistically significant. For our pleasantness linear mixed-effects model, we found statistically significant differences in the pairwise comparison between pre-satiety devalued ratings and post-satiety devalued ratings (estimate = -2.81, t-ratio = -6.55, SE = 0.42, p < 0.001) and in our pairwise comparisons of post-satiety devalued ratings and post-satiety non-devalued ratings (estimate = -1.78, t-ratio = -4.15, SE = 0.42, p = 0.0004). We did not find any significant differences between the pairwise comparisons of pre-satiety non-devalued ratings and pre-satiety devalued ratings (estimate = 0.15, t-ratio = 0.36, SE = 0.42, p = 0.983) nor with the pairwise comparisons of pre-satiety non-devalued ratings and post-satiety non-devalued ratings (estimate = -0.87, t-ratio = -2.04, SE = 0.42, p = 0.180). For our desirability linear mixed-effects model, we found statistically significant differences in our pairwise comparisons between pre-satiety non-devalued ratings and post-satiety non-devalued ratings (estimate = 2.25, t-ratio = -5.19, SE = 0.45, p < 0.001), in our pairwise comparisons between pre-satiety devalued ratings and post-satiety devalued ratings (estimate = 5.31, t-ratio = -11.59, SE = 0.45, p < 0.001), and in our pairwise comparisons of post-satiety non-devalued ratings and post-satiety devalued ratings (estimate = -2.93, t-ratio = -6.41, SE = 0.45, p < 0.001). We did not find any significant differences in our pairwise comparisons between pre-satiety non-devalued ratings and pre-satiety devalued ratings (estimate = 0.12, t-ratio = 0.27, SE = 0.45, p = 0.992).

Overall, the results from our behavioral analyses replicates previous human devaluation studies that also found participants reporting the devalued food as being affectively neutral as opposed to aversive after devaluation compared to before (Holland, 2008; Howard & Kahnt, 2017; Pool et al, 2019). While desirability ratings for both foods significantly decreased post-satiety compared to pre-satiety, the reduction in self-reported desirability was significantly more pronounced for devalued food compared to non-devalued food. Therefore, our results suggest that participants reported a change in value in the food that was specific to the one they had just eaten compared to the one they did not eat, despite finding both foods equally enjoyable and desirable prior to eating.

### Adjustments in behavioral strategies as a function of current motivational state

We first examined what kind of behavioral strategies influences the likelihood of switching responses by modeling the main and interaction effects of condition and feedback type with a binomial logistic regression with no reward pre-satiety as our reference point. In our model, we found significant differences in the main effects for devalued feedback (z = -10.73, coefficient = -0.66, SE = 0.06, p < 0.001 [CI 95%: -0.78, -0.53]), non-devalued feedback (z = -6.57, coefficient = -0.41, SE = 0.06, p < 0.001 [CI 95%: -0.54, -0.29]), and condition (z = -5.02, coefficient = -0.26, SE = 0.05, p < 0.001 [CI 95%: -0.37, -0.16]). Additionally, we found significant differences in the interaction effect for devalued feedback and condition (z = 3.48, coefficient = 0.29, SE = 0.08, p < 0.001 [CI 95%: 0.13, 0.47]) but we did not find significant differences in the interaction effect for non-devalued feedback and condition (z = -0.89, coefficient = -0.07, SE = 0.08, p = 0.373 [CI 95%: -0.25, 0.09]).

We then examined main effects of condition and feedback with a binomial logistic regression through two separate models with either pre-satiety devalued feedback or pre-satiety non-devalued feedback as our reference points. This approach allowed us to directly compare pre-satiety rewards and their respective switch rates for the post-satiety session. For our model with pre-satiety devalued feedback as our reference point, we found significant differences in the main effect for pre-satiety non-devalued feedback (z = 3.536, coefficient = 0.24, SE = 0.06, p = 0.656 [CI 95%: 0.10, 0.37]) but we did not find significant differences in the main effect for condition (z = -5.02, coefficient = -0.26, SE = 0.05, p < 0.001 [CI 95%: -0.37, -0.16]). For our model with pre-satiety non-devalued feedback as our reference point, we found significant differences in the main effect for condition (z = -5.09, coefficient -0.34, p < 0.001, [CI 95%: -0.47, -0.21]). While these results suggest differences in baseline switch rates between non-devalued feedback and devalued feedback, only the likelihood of switching responses significantly decreased for non-devalued feedback but not for devalued feedback.

Finally, we fitted the values of our model’s predictors by calculating the probability of the likelihood of switching (Figure 3). During the pre-satiety session, the probability of participants switching responses on the subsequent trial after receiving no reward feedback was 0.807 (or 80.7%), the probability of participants switching responses on the subsequent trial after receiving devalued feedback was 0.684 (or 68.4%), and the probability of participants switching responses on the subsequent trial after receiving non-devalued feedback was 0.734 (or 73.4%). During the post-satiety session, the probability of participants switching responses on the subsequent trial after receiving no reward feedback was 0.762 (or 76.2%), the probability of participants switching responses on the subsequent trial after receiving devalued feedback was 0.690 (or 69%), and the probability of participants switching responses on the subsequent trial after receiving non-devalued feedback was 0.661 (or 66.1%).

**Figure 3.**
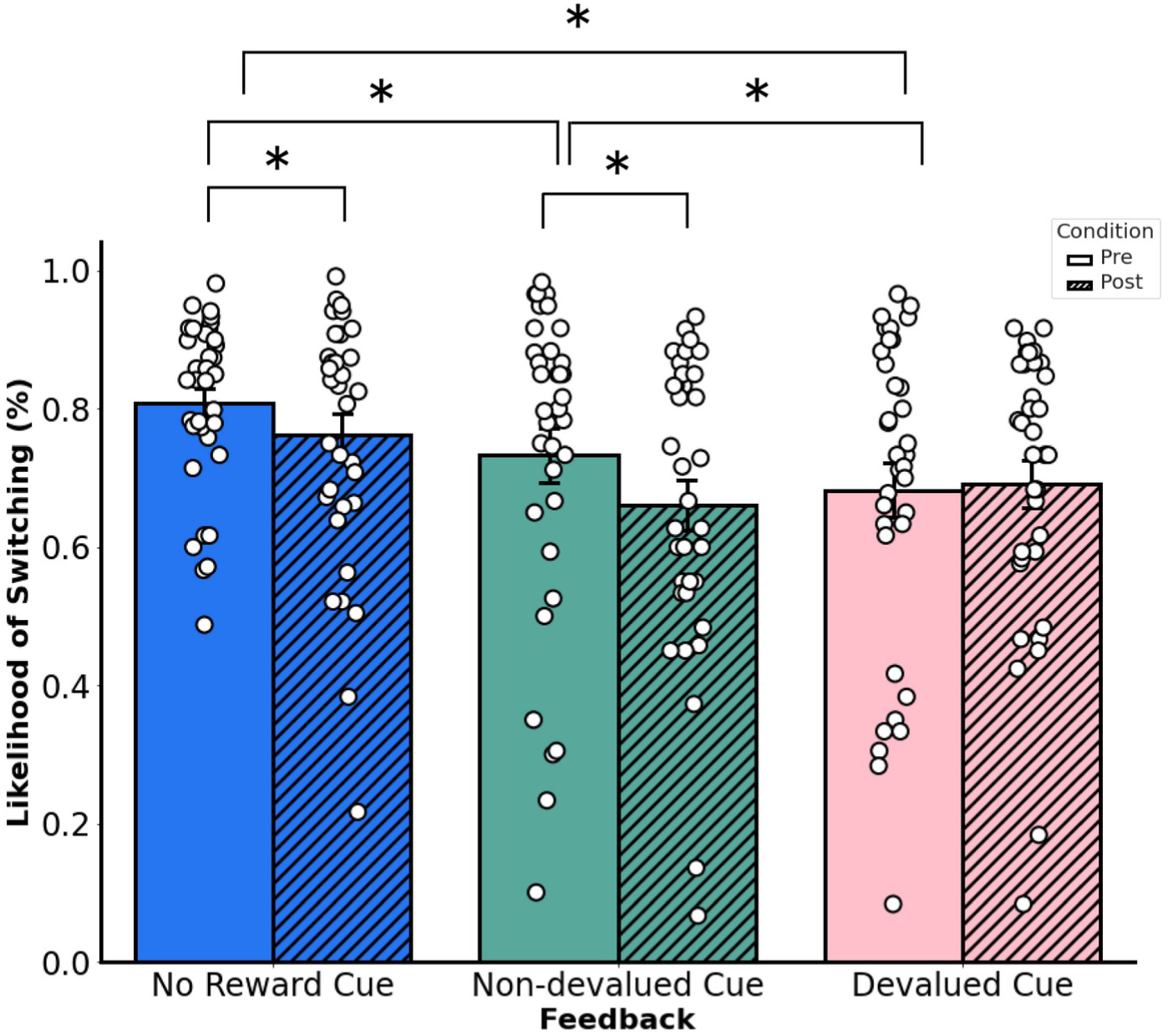
Average likelihood of switching door choice responses from the previous trial to the current trial based on feedback type. Changes between post- and pre-satiety sessions. Error bars denote the standard error of the mean.

The results suggest that participants were more likely to choose the same door on the subsequent trial similarly to the choice they made on the previous trial if they received reward feedback compared to no reward feedback during the pre-satiety session. During the post-satiety session, however, participants were more likely to choose the same door on the subsequent trial similarly to the choice they made on the previous trial if they received non-devalued feedback or no reward feedback; no significant adjustments in strategy were observed for devalued feedback between pre-satiety and post-satiety sessions. Together, this would suggest that participants made adjustments in their strategy in an outcome-specific manner even in a task where a learnable, optimal strategy was not possible. If participants were able to make adjustments in an outcome-specific manner, then this would also imply that participants were guided by an internal representation of goals reflective of the participant’s current motivational state.

### Non-parametric permutation tests examining ERP amplitudes reveal no significant differences following reinforcer devaluation

We first examined whether our novel Doors Tasks reliably evoked a RewP for either reward condition during the pre-satiety session. Figure 4 shows the spatiotemporal distribution for the feedback waveforms measured at channel Cz during pre-satiety (Figure 4A) and post-satiety recording sessions (Figure 4B). Figure 5 shows the spatiotemporal distribution for the Nondev RewP and Deval RewP measured at Cz calculated as the difference between wins and losses during pre-satiety (Figure 5A) and post-satiety recording sessions (Figure 5B). Figure 6 shows the grand averaged amplitudes for the Nondev RewP and Deval RewP at each time window of 189-239 ms, 240-290 ms, and 291-341 ms. In order to examine whether our task evoked a RewP, we performed two one-sample t-tests with our non-parametric permutations approach: 1) comparing the pre-satiety Nondev RewP against zero, and 2) comparing the pre-satiety Deval RewP against zero. Each type of one-samples t-test was performed at channels Fz, Cz, and Pz across our time windows of 189-239 ms, 240-290 ms, and 291-341 ms. When comparing the Nondev RewP against zero, we found significant differences for channel Fz at 240-290 ms (t = 4.38, maxT = 1.33, pcorr < 0.001), Fz at 291-341 ms (t = 2.49, maxT = 1.15, pcorr = 0.013), Cz at 240-290 ms (t = 5.13, maxT = 2.01, pcorr < 0.001), Cz at 291-341 ms (t = 3.19, maxT = 0.40, pcorr = 0.004), Pz at 240-290 ms (t = 6.90, maxT = 1.45, pcorr < 0.001), and Pz at 291-341 (t = 5.46, maxT = 0.03, pcorr < 0.001); we did not find significant differences for channel Fz at 189-239 ms (t = 1.31, maxT = 1.49, pcorr = 0.180), Cz at 189-239 ms (t = 1.39, maxT = 1.40, pcorr = 0.153), or for channel Pz at 189-239 ms (t = 0.99, maxT = 1.57, pcorr = 0.320). When comparing the Deval RewP against zero, we found significant differences for channel Fz at 240-290 ms (t = 2.41, maxT = 1.42, pcorr = 0.020), Cz at 240-290 ms (t = 3.60, maxT = 0.59, pcorr < 0.001), Pz at 240-290 ms (t = 6.09, maxT = 0.01, pcorr < 0.001), and Pz at 289-341 ms (t = 4.33, maxT = 0.85, pcorr < 0.001); we did not find significant differences for channel Fz at 189-239 ms (t = 0.48, maxT = 1.07, pcorr = 0.628), Fz at 291-341 ms (t = 0.42, maxT = 1.57, pcorr = 0.657), Cz at 189-239 ms (t = 1.23, maxT = 1.28, pcorr = 0.260), Cz at 291-341 (t = 1.09, maxT = 1.56, pcorr = 0.292), or for channel Pz at 189-239 ms (t = 0.81, maxT = 1.48, pcorr = 0.418). Our findings suggest that our task was able to reliably evoke a RewP for either reward condition during the pre-satiety session across comparable time windows and channel sites as described in prior RewP studies (Proudfit, 2015; Krigolson, 2018).

**Figure 4.**
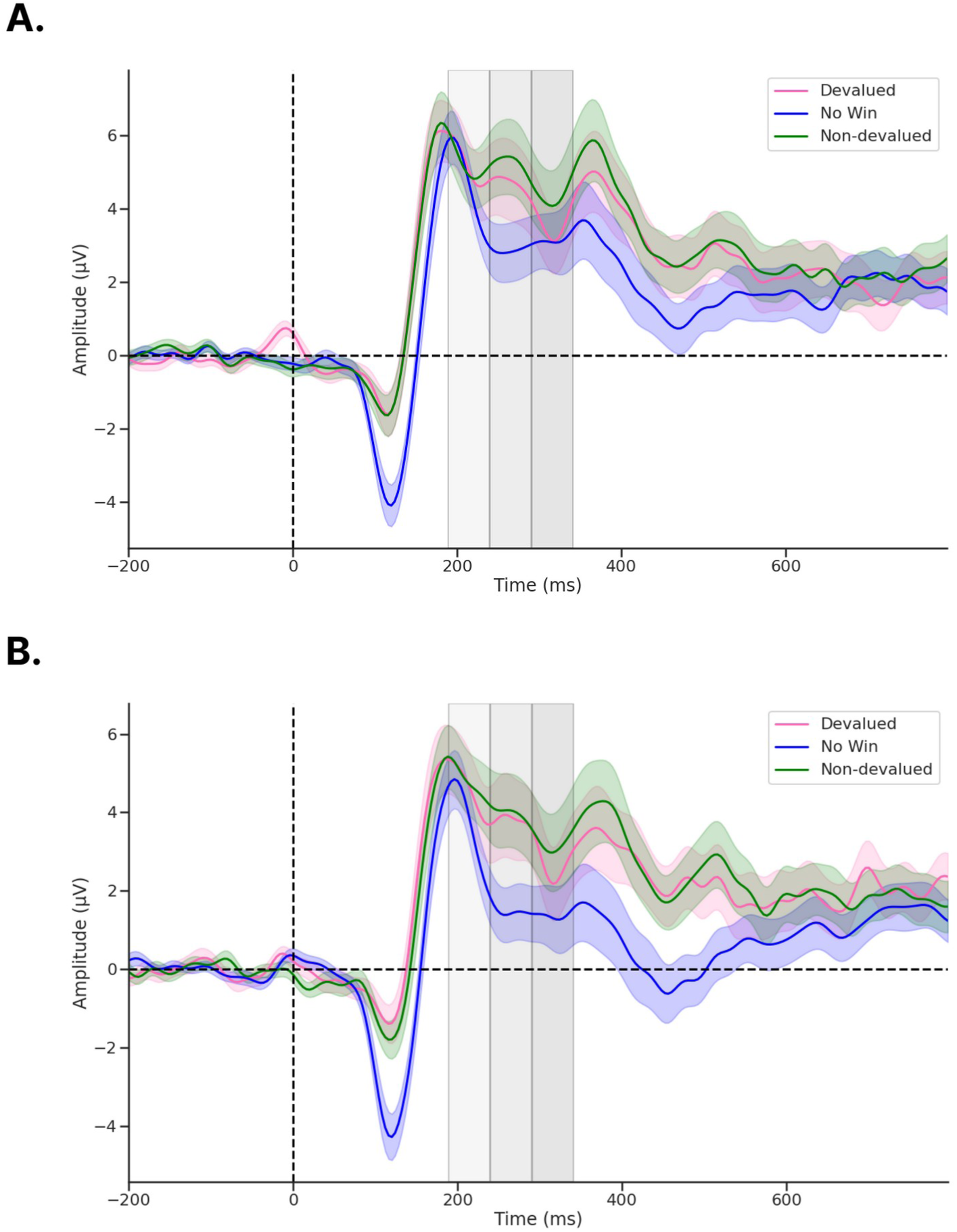
ERPs recorded from channel Cz following feedback onset. a) During pre-devaluation, and b) during post-devaluation. Each gray shaded region denotes time windows 189-239 ms (light gray), 240-290 ms (medium gray), and 291-341 ms (dark gray). ERP shading corresponds to the standard error (SE).

**Figure 5.**
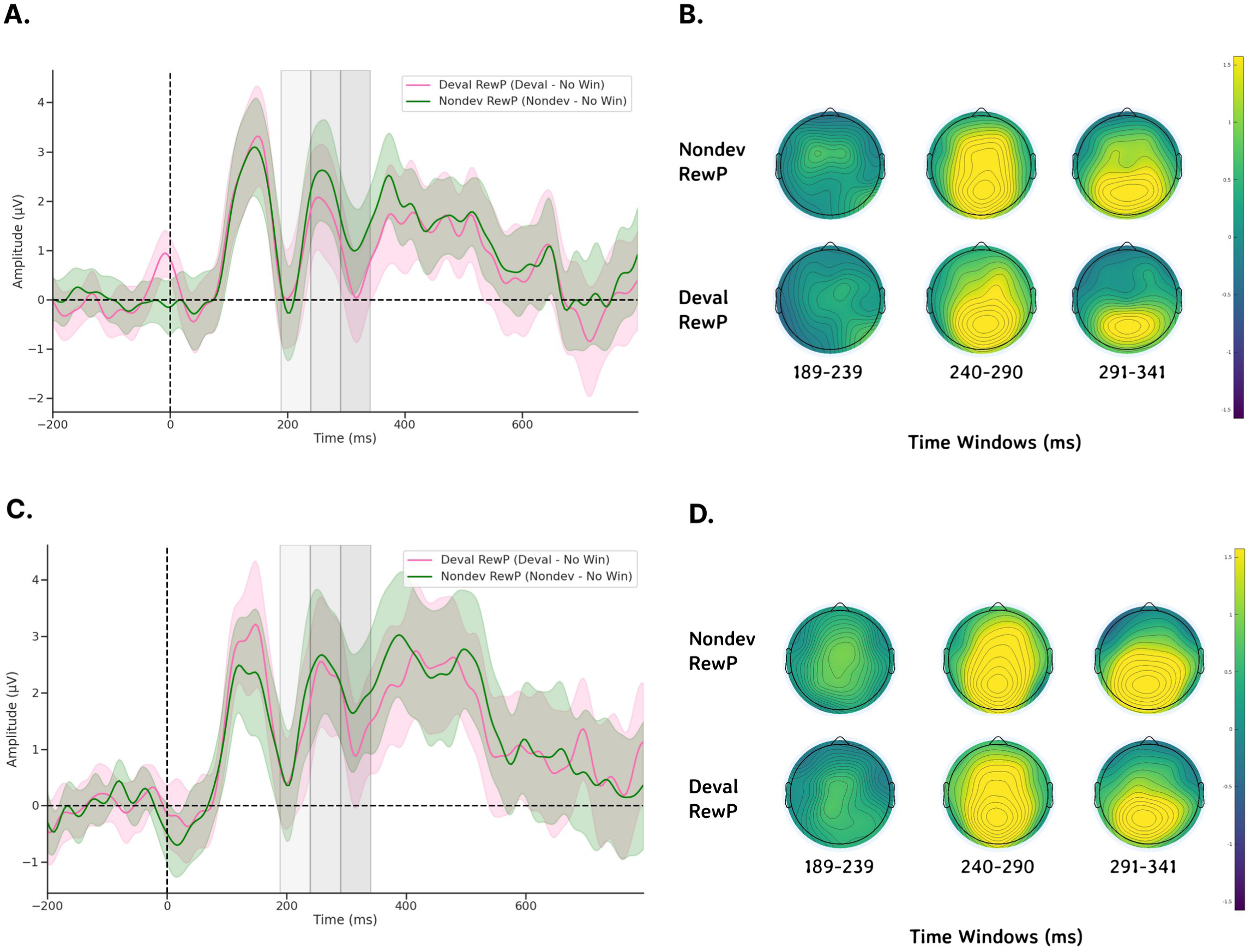
Topological maps and difference waveforms of non-devalued and devalued feedback recorded from channel Cz following feedback onset. a) Pre-devaluation waveforms at Cz, b) pre-devaluation topological maps for Nondev and Deval RewPs at each time window, c) post-devaluation waveforms, and d) post-devaluation topological maps for Nondev and Deval RewPs at each time window. Each gray shaded region denotes time windows 189-239 ms (light gray), 240-290 ms (medium gray), and 291-341 ms (dark gray). ERP shading corresponds to 95% confidence intervals (CI).

**Figure 6.**
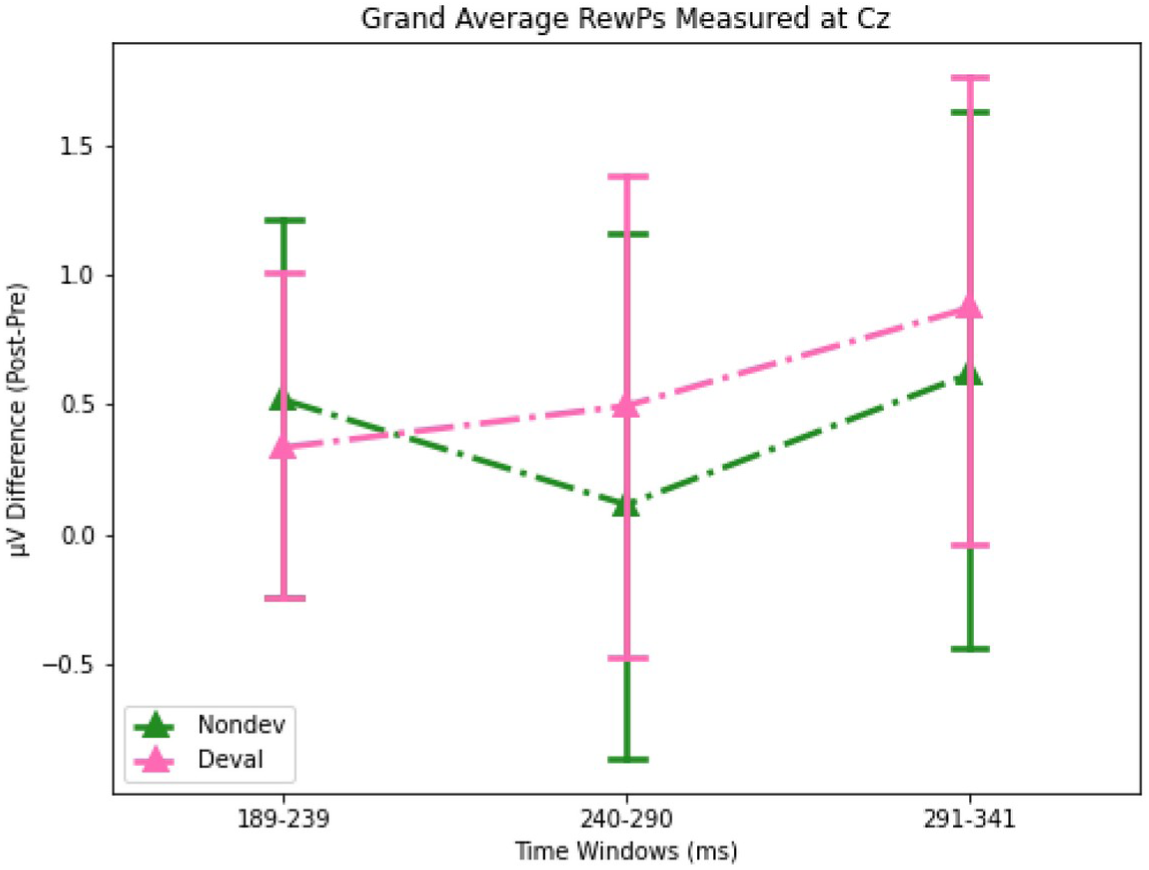
Barplots for grand averaged (post-pre) amplitudes for Nondev and Deval RewPs measured at Cz for each time window. Error bars correspond to 95% confidence intervals (CI).

We then examined whether participants responded to non-devalued feedback and devalued feedback across both sessions by testing the interaction effects between reward type and session type. We compared the grand average Nondev RewP with the grand average Deval RewP at our time window of 189-239 ms (Figure 4A & 4B). For this analysis, we found no significant differences in Fz (t = 0.15, maxT = 2.15, pcorr = 0.883), Cz (t = 0.41, maxT = 1.92, pcorr = 0.691), or Pz (t = 0.56, maxT = 2.10, pcorr = 0.584) within the time window of 189-239 ms post-feedback. Next, we compared the grand average Nondev RewP with the grand average Deval RewP at our time window of 240-290 ms. For this analysis, we found no significant differences in Fz (t = 1.26, maxT = 1.99, pcorr = 0.216), Cz (t = 0.70, maxT = 2.06, pcorr = 0.483), or Pz (t = 0.07, maxT = 1.98, pcorr = 0.937) within the time window of 240-290 ms post-feedback. Finally, we compared the grand average Nondev RewP with the grand average Deval RewP at our time window of 291-341 ms, corresponding to where we identified the most negative trough measured for pre-satiety reward feedback between 240-340 ms (Sambrook & Goslin, 2015). For this analysis, we did not find any significant differences in Fz (t = 1.39, maxT = 1.99, pcorr = 0.170), Cz (t = 0.44, maxT = 2.07, pcorr = 0.655), or Pz (t = 0.46, maxT = 2.02, pcorr = 0.609) within the time window of 291-341 ms post-feedback.

In order to directly examine interaction effects between food type and session, we calculated the difference between non-devalued feedback and devalued feedback, which we refer to as our Reward Contrast. We then performed a one-sample t-test with our non-parametric permutations approach to examine if the grand average of our Reward Contrast (Post Reward Contrast – Pre Reward Contrast) was significantly greater than zero. Our null hypothesis assumes no differences between devalued and non-devalued feedback regardless of session (pre vs post). When comparing the grand average Reward Contrast against zero, we did not find any significant differences at channel Fz at 189-239 ms (t = 0.93, maxT = 0.611, pcorr = 0.368), Fz at 240-290 ms (t = 0.40, maxT = 1.07, pcorr = 0.689), Fz at 291-341 ms (t = 1.39, maxT = 2.06, pcorr = 0.174), Cz at 189-239 ms (t = 1.70, maxT = 1.72, pcorr = 0.090), Cz at 240-290 ms (t = 1.68, maxT = 1.89, pcorr = 0.088), Cz at 291-341 ms (t = 0.44, maxT = 3.03, pcorr = 0.667), Pz at 189-239 ms (t = 0.78, maxT = 1.74, pcorr = 0.433), Pz at 240-290 ms (t = 1.81, maxT = 2.50, pcorr = 0.085), or Pz at 291-341 ms (t = 0.46, maxT = 2.42, pcorr = 0.670) post-feedback.

Finally, we investigated whether the mean amplitude for the P1 was greater for devalued feedback compared to non-devalued feedback following devaluation as previously reported (Luque et al., 2017). For this analysis, we did not find any significant differences in Oz (t = 1.41, maxT = 2.10, pcorr = 0.164) within the time window 101-151 ms post-feedback. Together, this suggests that neither reward feedback nor the RewP differed in amplitudes regardless of whether this was before or after reinforcer devaluation, suggesting an insensitivity to changes in current reward value.

## Discussion

To examine the hypothesis that the RewP is sensitive to reward value as a function of the current motivational state, we compared RewP responses involving two distinct reinforcers before and after reinforcer devaluation. In contrast to our behavioral results and main hypothesis, we found that the mean amplitude of the Nondev and Deval RewPs did not significantly differ before or after satiety. Our RewP amplitude results do not appear to be explained by novelty effects since rewards and non-rewards were equally probable, re-learning associations in a diminished state, disproportionate pleasantness ratings at pre-satiety, or visual features of the feedback. Since pre-satiety recordings of devalued and non-devalued feedback were similar, this would support previous research suggesting the RewP is sensitive to subjective preferences of rewards (Brown et al., 2022; Peterburs et al., 2019). However, our null results suggest prior studies might have shown that the RewP signal general properties of reward as opposed to its sensory-specific properties.

Our lack of devaluation effects is unlikely due to the possibility of missing the RewP outside of our pre-defined time window. Here, we defined our time window *a priori* from a meta-analysis suggesting the RewP is maximal at any time point between 240-340 ms post-feedback (Sambrook & Goslin, 2015). We then further narrowed down this time window using a collapsed localizer approach based on the latency of the N2 for reward feedback, resulting in a time-window of 291-341 ms (Luck & Gaspelin, 2017). In addition to analyzing a time window centered around the latency of the N2, we also analyzed two preceding 50 ms time windows—189-239 ms and 240-290 ms—since the RewP can vary considerably in its latency (Sambrook & Goslin, 2015). With these three time windows, we first examined whether RewP amplitudes recorded during the pre-satiety session significantly differed from zero. We found a positivity maximal at 240-290 ms and with a centroparietal distribution. Although this positivity was found earlier than our initial time window of 291-341 ms, the RewP being maximal at 240-290 ms aligns with prior reports regarding the latency of the RewP (Sambrook & Goslin, 2015). While this positivity being maximal with a centroparietal distribution is a slight departure from canonical descriptions of the RewP having a frontocentral topography, a RewP with a centroparietal topography has been observed in other studies (Cavanagh, 2015; Brown et al., 2021). Moreover, the results from our one-samples t-test showed that we reliably evoked a RewP for non-devalued and devalued feedback at channels Fz, Cz, and Pz in the 240-290 ms time-window. When we tested for interaction effects of our grand averaged RewP, however, we found no significant differences across all channels and time-windows tested. Therefore, it’s unlikely that we did not find any effects due to having missed the RewP because we measured different channels and time windows as informed by the literature.

It is also unlikely our findings that the RewP lacking sensitivity toward devaluation reflects a contribution of habitual responding to the generation of the RewP. First, participants’ performances on the self-reported pleasantness scales indicate a selective shift in perceived pleasantness of the food eaten to satiety compared to the food that wasn’t eaten. The influence of satiety toward participants’ ratings of pleasantness supports previous human devaluation studies detecting the presence of the devaluation effect (Rolls et al., 1981; Rolls et al., 1983; Gottfried et al., 2003; Howard & Kahnt, 2017; Reber et al., 2017; Pool et al., 2019; Thompson et al., pre-print). Second, the results from our exploratory switch-stay analysis shows that the likelihood of switching door responses upon receiving non-devalued reward feedback decreased during post-satiety compared to pre-satiety, whereas the likelihood of switching door responses upon receiving devalued feedback remained unchanged. Habitual behavior has been referred to as a function of repetitive behavior and reflects stimuli exerting more control over behavior than outcome values (Tricomi et al., 2009). If the behavior was habitual, then participants would either continue behavior at levels similar prior to devaluation or increasing responding toward devalued outcomes following devaluation because participants were immune to changes in reward value (Tricomi et al., 2009; Reber et al., 2017). We did not find evidence of either type of behavior from participants. Rather, since participants were using the identity of the reward to make adjustments in making choices on future trials, this would instead suggest participants were using an internal model to prospectively guide behavior (Wilson et al., 2014). Third, habitual behavior is notoriously difficult to induce in human participants and likely reflects the preservation of value-free learning about repetitive actions, to which we did not observe from our participants (de Wit et al., 2018; Miller et al., 2019; Pool et al., 2021; Nebe et al., 2024). Finally, not all devaluation-insensitive behavior can be considered habitual as it can be alternatively explained by knowledge retention (Pool et al., 2019), incomplete knowledge transfer (Reber et al., 2017), variations in experimental parameters (Holland, 2008; Tricomi et al., 2009; Klossek & Dickinson, 2011), or the effectiveness of the reinforcer devaluation procedure based on the type of reinforcer used (Seabrooke et al., 2017).

We also sought to replicate findings previously reported on the early visual component, the P1, demonstrating increased amplitudes recorded from occipital electrodes toward devalued outcomes, suggesting devaluation-insensitive signals important for perceptual prioritization relevant for behavioral adaptation (Luque et al., 2017). Unlike the previous study, we fail to find any significant differences in mean P1 amplitudes between devalued and non-devalued feedback between pre-satiety and post-satiety sessions. There are a couple of interpretations for why we failed to replicate findings from Luque and colleagues (2017). First, the devaluation technique used in Luque et al. (2017) was verbal instruction, whereas this study used satiety. Although verbal instructions have been used as an effective method of reinforcer devaluation (Allman et al., 2010; Thompson et al., pre-print), it is unclear whether devaluation-insensitive P1 amplitudes are unique to cognitive forms of devaluation versus those that manipulate motivational or physiological states. Second, although Luque and colleagues (2017) had associated cues with outcomes, the outcomes pertained to various magnitudes of one reinforcer. Learning the association involving a single reinforcer versus several reinforcers are more likely to promote the formation of habits because the identity of reinforcers become increasingly irrelevant as training progresses (Klossek et al., 2008; Tricomi et al., 2009; Klossek & Dickinson, 2011). Future studies could compare various forms of devaluation affects P1 amplitudes by learning a single action-outcome contingency versus two action-outcome contingencies per trial in conjunction with response rates toward devalued and non-devalued reinforcers.

One explanation parsimonious to previous work and this study’s findings is that the RewP could reflect knowledge retention of task-relevant features of the reward. For our task, we employed Pavlovian conditioning between our feedback (i.e., cues) and pictures of outcomes during the pre-satiety session and presenting only the cues in extinction during the post-satiety session. In previous devaluation tasks using Pavlovian learning, some sensory-specific features of representations such as its identity can be sensitive to devaluation whereas other sensory-specific features such as its spatial location were insensitive to devaluation, suggesting that some forms of task-relevant information are malleable to change whereas other forms of information are held constant (Pool et al., 2019; Pool et al., 2021). Here, it’s possible that the devaluation insensitivity we’ve observed with RewP amplitudes could reflect a devaluation-insensitive retention of sensory-specific features of the reinforcer. If knowledge retention of the sensory-specific properties were represented, then this could also explain why previous RewP studies found modulations of current reward value while we did not: if feedback about rewarding outcomes were directly experienced instead of inferring them, then it’s possible that participants were re-learning associations about rewards but in a different state. Banica and colleagues (2023) found hungry participants showed greater RewP amplitudes for food rewards compared to monetary rewards, whereas sated participants showed similar RewP amplitudes for food and monetary rewards. In Banica et al. (2023), however, the delivery of this feedback was not inferred and directly experienced instead. Therefore, the participants might not need to retain certain types of knowledge about the outcomes they received because participants were continuously learning about reward feedback.

Alternatively, it’s possible that a devaluation insensitive RewP reflects reward value being better or worse than expected as a function of the task type. The Doors Task, which is commonly used to evoke a RewP, measures how participants respond to positive or negative feedback based on performance using guessing (Proudfit, 2015). In a guessing task, participants learn about their performance across time only upon receiving feedback, whereas tasks utilizing planning strategies involve learning about performance through rule sets, contingencies, and outcomes (Elliot et al., 1998). Although our participants inferred the associations between feedback and reinforcers, they were unable to infer sensory-specific information about reinforcers at the time of choice. Rather, the only information our participants could infer at the time of choice is if they will receive a food reward or no reward because there was no information to infer about the receipt of one type of food reward being more likely than another. This could explain why previous work found contextual sensitivity in the RewP while we did not. For example, winning tokens for a casino with a higher payout evoked a larger RewP than winning tokens for a casino with a lower payout because participants were able to see different games within each casino and learn their overall payout rates (Umemoto et al., 2017).

Additionally, a modified Doors Tasks requiring to learn associations between colored doors and shock expectancy evoked a RewP in participants when receiving unexpected feedback about avoiding shocks (Bauer et al., 2024). These findings would suggest information provided at the time of choice influenced whether feedback was better or worse than expected and this influence was potentiated by the unexpectedness of the type of feedback. An interesting prospect for future work could examine whether providing an environment in which inferred sensory-specific information about a food reinforcer at the time of choice could modulate the RewP following devaluation and whether surprise contributes toward an additive effect on perceiving feedback.

Finally, future work could examine whether the RewP is devaluation-insensitive when decreasing the value of secondary reinforcers such as money or social rewards. It’s unclear whether devaluation-insensitive RewPs toward representations of food reward reflects domain-specific or domain-general processing of reward representations. Investigating this could be possible by utilizing the task described here and protocols which have robustly measured devaluation effects toward money (Allman et al., 2010), social rewards in the form of smiling faces (Thompson et al., pre-print), and video clips from children’s cartoons (Klossek et al., 2008). Prior work has suggested overlapping yet dissociable networks in how we perceive different types of rewards. For example, food deprivation and social deprivation activate similar patterns of midbrain activity in humans when viewing pictures of food or social rewards during states of deprivation (Tomova et al., 2020). In non-human primates, however, ablating amygdala connectivity with orbital networks of the prefrontal cortex impaired food devaluation but ablating amygdala connectivity with medial networks of the prefrontal cortex impaired social devaluation (Pujara et al., 2022). Experiments successfully devaluing other rewards, along with experiments systematically requiring a fasting of food and social rewards, matching rewards on pleasantness and desirability, and devaluing those rewards could yield further insight into what kind of precise mechanisms contribute to the RewP.

There are several limitations to this study worth noting for future inquiry. First, a limitation with our study is that omitting the outcome pictures during post-satiety could have resulted in devaluation-insensitive RewPs. Since testing how participants inferred current reward value required feedback presented during extinction, presenting the outcomes during post-satiety would have introduced a key confound. We believe future work could replicate this study without omitting the outcome screen during the post-satiety session. Another limitation is it’s unclear why participants used Pavlovian feedback as a heuristic strategy during our Doors Task and how this behavior relates to more established measures of outcome expectancy indexing Pavlovian conditioning (e.g., pupil dilation). A future experiment combining our version of the Doors Task with measures of Pavlovian conditioning such as pupillometry could address how door switching phenomena relates to Pavlovian expectations (Pool et al., 2019).

Together, this study is the first to demonstrate that an ERP implicated in reward processing is insensitive to reinforcer devaluation despite participants endorsing devaluation-sensitive behavior. The current findings contradict recent proposals suggesting the RewP encodes reward value relevant to current motivational state (Baker et al., 2016; Peterburs et al., 2019; Banica et al., 2023; Brown et al., 2022; Proudfit, 2015). These results suggest that although the RewP is sensitive to reward liking and demonstrates contextual sensitivity, factors pertinent to reward value might provide different contributions toward the RewP depending upon whether those factors were inferred. Utilizing paradigms such as those from behavioral neuroscience and reinforcement learning could refine understanding the computational processes represented within the RewP and, broadly, provide a more comprehensive account on the neural contributors of goal-directed behavior.

## Declarations

### Ethics approval

The study was approved by the George Mason University International Review Board and in accordance with guidelines from the 1964 Declaration of Helsinki.

### Consent to participate

Informed consent was obtained from all individual participants included in the study.

### Consent to publication

Not applicable.

### Availability of data and materials

Data or materials for the experiment are available upon request to the corresponding author and the experiment was not preregistered.

### Code availability

Custom code for the experiment are available upon request to the corresponding author.

### Authors’ contributions

**Lindsay S. Shaffer:** Conceptualization; methodology; data collection; analysis; writing – original draft, reviewing, and editing. **Holly D. Crowder:** Data collection; analysis. **Peter A. Kakalec:** Analysis; writing – editing. **Lam T. Duong:** Methodology; data collection. **Craig G. McDonald:** Conceptualization; methodology; analysis; writing – reviewing and editing. **James C. Thompson:** Conceptualization; methodology; analysis; writing – reviewing and editing; supervision.

## Acknowledgments

We thank Ms. Allison Haddad, Ms. Rebecca Roy, and Ms. Alexa Young for their help with recruitment and Dr. James Cavanagh, Dr. Cameron Hassall, Ms. Eslam Hassan, Dr. Benjamin Hayden, and Dr. Daniel Roberts for helpful discussions. We also would like to thank our editor and reviewers for their insightful comments during the review process.

