## Supplementary Material for "The reward positivity is insensitive to reinforcer devaluation"

***Changes in RewP amplitudes not correlated with changes in self-report measures or amount of consumption.*** We performed a Pearson's correlation coefficient to assess the linear relationship between individual valuation changes in pleasantness and changes in our RewP amplitudes measured at our initial time window of 291-341 ms. For individual valuation changes, we calculated a difference score between post vs pre in pleasantness each for the devalued and non-devalued reinforcer. For changes in RewP amplitudes, we calculated a grand average difference (Post-satiety RewP – Pre-Satiety RewP). Since not all of our EEG participants (n=31) had self-reported measures, the final sample size for this analysis is 27 participants. When comparing changes in non-devalued RewP amplitudes with changes in pleasantness ratings for non-devalued food, we did not find significant differences (coefficient = 0.028, p = 0.886). When comparing changes in devalued RewP amplitudes with changes in pleasantness ratings for devalued food, we did not find significant differences (coefficient = -0.137, p = 0.494).

We then performed a Pearson's correlation coefficient to assess the linear relationship between individual valuation changes in desirability and changes in our RewP amplitude measured at our initial time window of 291-341 ms (n=27). For individual valuation changes, we calculated a difference score between post vs pre in desirability each for the devalued and non-devalued reinforcer. When comparing changes in non-devalued RewP amplitudes with changes in desirability ratings for non-devalued food, we did not find significant differences (coefficient = 0.050, p = 0.803). When comparing changes in devalued RewP amplitudes with changes in desirability ratings for devalued food, we did not find significant differences (coefficient = 0.295, p = 0.135).

Finally, we performed a Pearson's correlation coefficient to assess the linear relationship between consumption of food or water and changes in either our non-devalued RewP or devalued RewP amplitude measured at our initial time window of 291-341 ms (n=29). When comparing changes in non-devalued RewP amplitudes, we did not found significant differences with food consumption (coefficient = 0.195, p = 0.310) or for water consumption (coefficient = 0.101, p = 0.598). When comparing changes in devalued RewP amplitudes, we did not found significant differences with food consumption (coefficient = 0.045, p = 0.812) or for water consumption (coefficient = 0.045, p = 0.814).

***Wilkinson notation for linear mixed-effects models.*** We provide below the syntax for each of our linear mixed-effects models used in this study (Wilkinson & Rogers, 1973).

For pleasantness, we defined our model as:

(1) *pleasantness ~ food type * session + ( ~1 | participant)*

For desirability, we defined our model as:

(2) *desirability ~ food type * session + ( ~1 | participant)*

For hunger, we defined our model as:

(3) *hunger ~ session + ( ~1 | participant)*

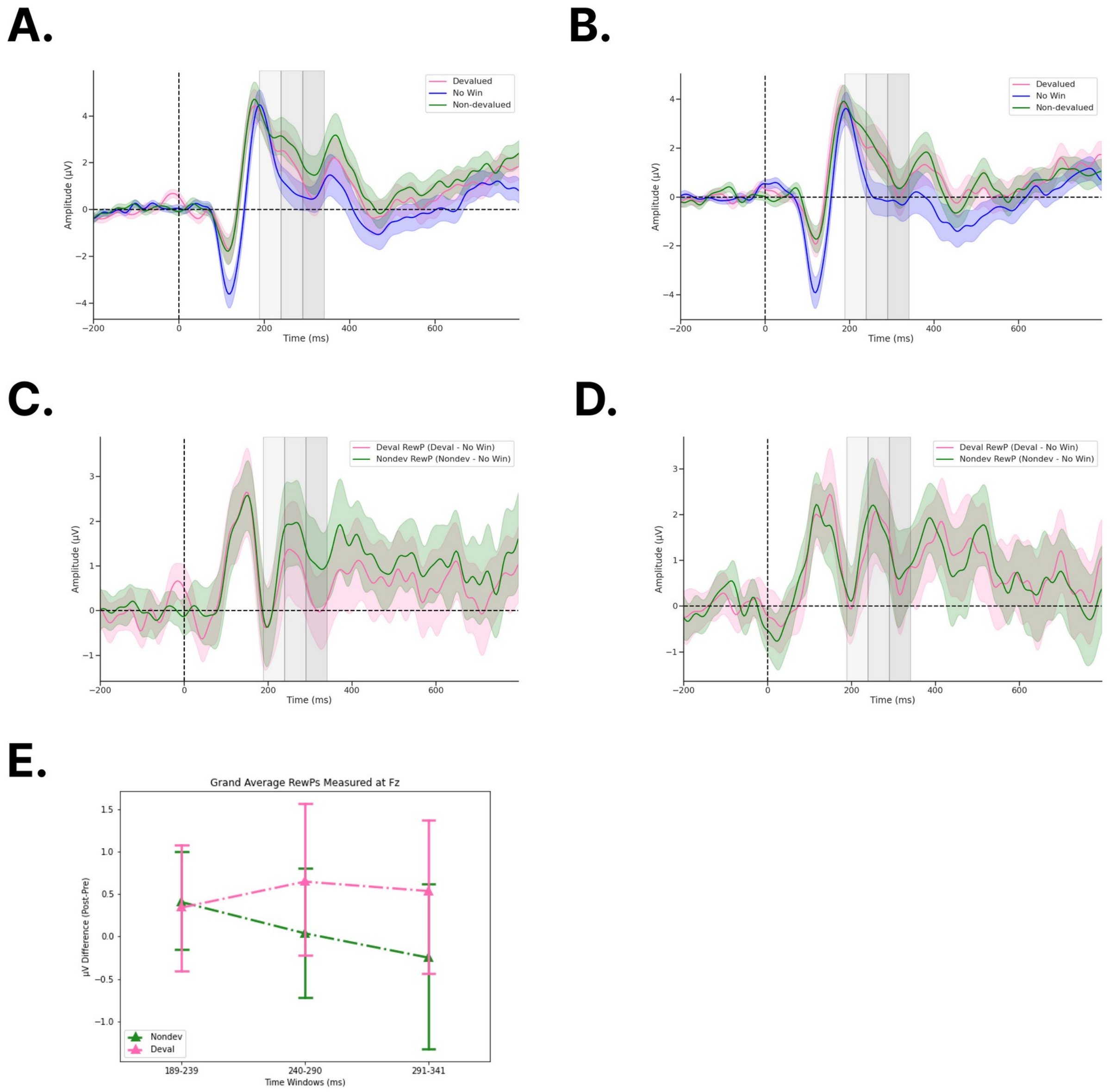


**Figure S1. ERPs recorded from channel Fz following feedback onset.** a) Raw feedback during pre-devaluation, b) raw feedback during post-devaluation, c) difference waveforms pre-devaluation, d) difference waveforms post-devaluation, and e) barplots for grand averaged (post-pre) amplitudes for Nondev and Deval RewPs measured at Fz for each time window. Each gray shaded region denotes time windows 189-239 ms (light gray), 240-290 ms (medium gray), and 291-341 ms (dark gray). ERP shading in S1A and S1B corresponds to the standard error (SE). ERP shading in S1C, S1D, and error bars in S1E correspond to 95% confidence intervals (CI).

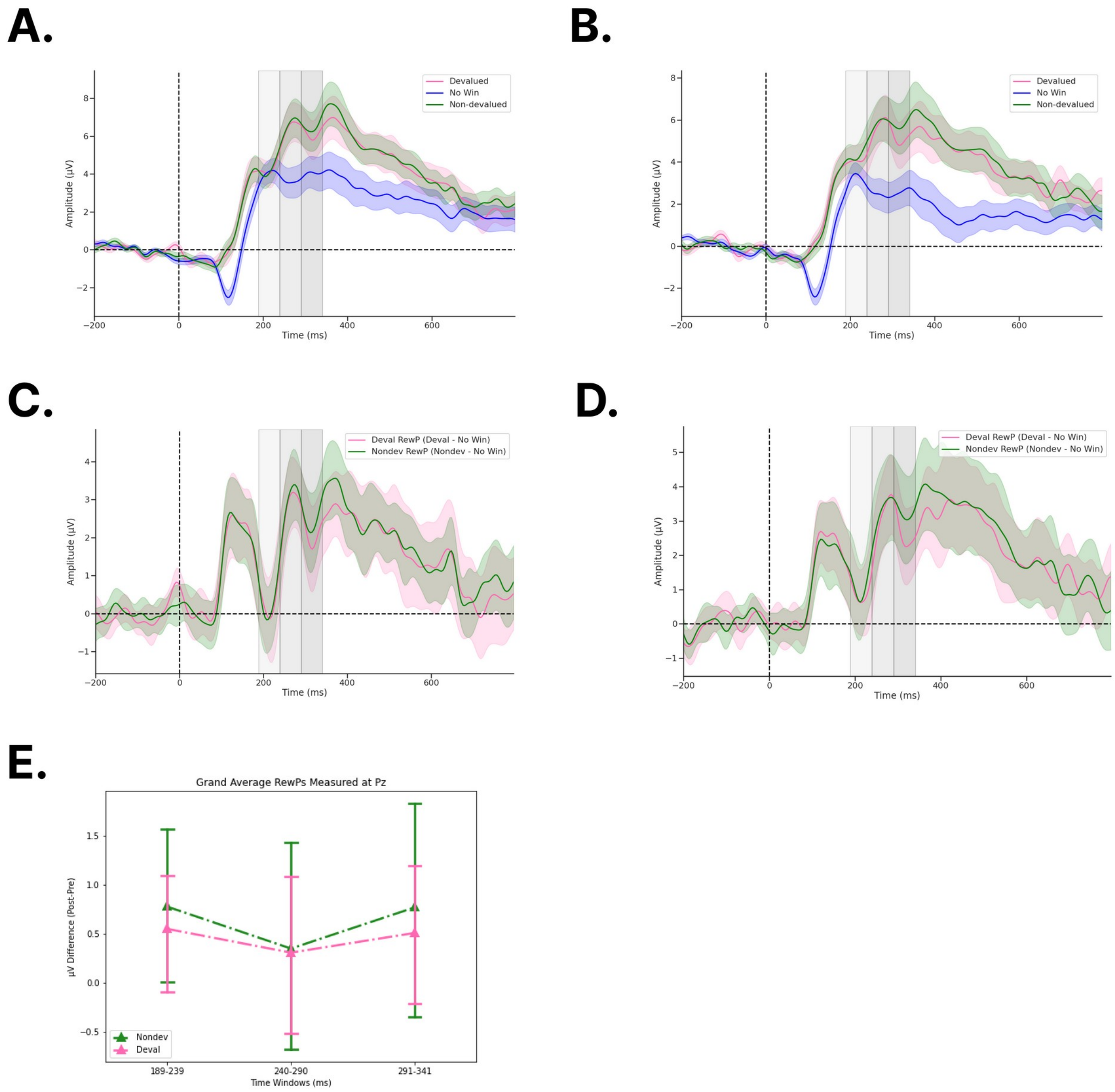


**Figure S2. ERPs recorded from channel Pz following feedback onset.** a) Raw feedback during pre-devaluation, b) raw feedback during post-devaluation, c) difference waveforms pre-devaluation, d) difference waveforms post-devaluation, and e) barplots for grand averaged (post-pre) amplitudes for Nondev and Deval RewPs measured at Pz for each time window. Each gray shaded region denotes time windows 189-239 ms (light gray), 240-290 ms (medium gray), and 291-341 ms (dark gray). ERP shading in S2A and S2B corresponds to the standard error (SE). ERP shading in S2C, S2D, and error bars in S2E correspond to 95% confidence intervals (CI).
